# Analysis of wild plant pathogen populations reveals a signal of adaptation in genes evolving for survival in agriculture in the beet rust pathogen (*Uromyces beticola*)

**DOI:** 10.1101/2021.08.12.456076

**Authors:** Mark M^c^Mullan, Lawrence Percival-Alwyn, Kevin Sawford, Gemy Kaithakottil, Michelle Grey, Hélène Yvanne, Ross Low, Sally Warring, Darren Heavens, Ned Peel, Jakob Kroboth, Mark Stevens, David Swarbreck, Matt Clark, Neil Hall

## Abstract

Improvements in crop resistance to pathogens can reduce yield losses and address global malnourishment today. Gene-for-gene -type interactions can identify new sources of resistance but genetic resistance is often short lived. Ultimately an understanding of how pathogens rapidly adapt will allow us to both increase resistance gene durability and more effectively target chemical treatments. Until recently all agricultural pathogens were living on wild hosts. To understand crop pathogen evolution, we compared genetic diversity in agricultural and wild populations. Wild reservoirs may be the source of emergent pathogen lineages, but here we outline a strategy for comparison of wild and agricultural pathogen populations to highlight genes adapting to agriculture. To address this, we have selected and developed the beet rust system (*Beta vulgaris*, *Uromyces beticola*, respectively) as our wild-agricultural model. Our hypothesis is that pathogen adaptation to agricultural crops will be evident as divergence in comparisons of wild and agricultural plant pathogen populations. We sampled isolates in both the wild and agriculture, sequenced and assembled and annotated a large fungal genome and analysed genetic diversity in 42 re-sequenced rust isolates. We found population differentiation between isolates in the wild compared to a predominantly agricultural group. Fungal effector genes are co-evolving with host resistance and are important for successful colonisation. We predicted (and found) that these exhibit a greater signal of diversification and adaptation and more importantly displayed increased wild agricultural divergence. Finding a signal of adaptation in these genes highlights this as an important strategy to identify genes which are key to pathogen success, that analysis of agricultural isolates alone cannot.

**Author Summary:** As quickly as we develop new strategies for crop defence, pathogens evolve to circumvent them. Novel crop pathogen strains emerge periodically and sweep through the agricultural system. However, because of the (often) clonal nature of these crop pathogens it is difficult to identify the trait that is key to their success. In other words, if there is a trait that is key for success in agriculture, all agricultural isolates will have it (or die without it). What we need is a case and control system where we identify genes important to pathogen success in agricultural by comparing them to pathogens that live in the wild. Here we exemplify this strategy by focussing on genes already known to specifically adapt for the successful colonisation of the host, the fungal effector genes. We find that these genes appear to be evolving quickly and that they are more different between the wild and agriculture than other non-effector genes. These differences between wild and agricultural pathogens suggest we are observing adaptation to agriculture. We do this work in the sugar beet rust system because of its tractability to sample but this understanding about how to identify genetic variation that is key to pathogen success in agriculture is applicable to crop systems where pathogen reservoirs exist as well as other pathogen reservoir systems (e.g. zoonoses).

## Introduction

By 2050 we must feed nine billion people, yet two billion people are currently malnourished and over 20% of agricultural crops are lost to disease annually (1,2). Crop protection by genetic resistance is characterised by short disease free periods, or a lag before pathogens adapt (3,4). Once resistance is broken, emergent diseases spread quickly through the host population and new sources of resistance are needed, which take several years to develop. Wild crop progenitor species are currently being explored as potential reservoirs of resistance gene diversity (3,5). However, wild host species may also be reservoirs for pathogens (6). On the one hand genetic variation present in these wild pathogens could be considered a source of novel genetic variation pre-adapted to the next round of resistance, on the other these wild pathogens represent a unique resource that can also be used to identify the characteristics important for pathogen success in agriculture. Wild and agricultural environments are different on a number of levels, primarily host genetic diversity and density but also in terms of other abiotic factors such as water availability, light levels and use of fertilisers and chemical control agents (7). These measures are expected to impact both host and pathogen directly. Analyses of emergent agricultural pathogens that omit genetic diversity present in pathogen populations from wild crops relatives, are not expected to identify the most important characteristics for success in agriculture because this critical polymorphism is likely fixed in agriculture. Instead, what is needed is a comparison of agricultural isolates to others that have not invaded (a case and a control).

Agriculture is a relatively recent phenomenon, as such, all agricultural pathogens recently lived on a wild host and, since the invention of agriculture, have either speciated, or are still exchanging genetic variation with wild counterparts. Pathogen reservoirs of genetic diversity represents a continuum from one-off hybridisation and introgression events, through to contemporary gene flow between populations living in different environments. Characterisation of one off hybridisation events has been implicated in the generation and success of plant pathogens such as Dutch Elm Disease (8) and *Zymoseptoria pseudotritici* (9) as well the human malaria pathogen (10). At an intermediate level, in pathogens such as *Albugo candida* (an oomycete which can infect over 200 plant species), each lineage apparently diverged from other host specific lineages with the exception of for rare recombination or introgression events between lineages (11,12). At the other end of the spectrum, invasion biology and population genetics has been considering the importance of genetic diversity, adaptive potential and invasion success (13). However, these processes are inherently difficult to study because they are rare, either they are a single hybrid speciation event, or they are one off invasion events (6,14). For example, the ash dieback pathogen (*Hymenoscyphus fraxineus*) has been a highly successful invader of Europe from Asia (15). Population genetic analyses applied to the invasive and source populations has shown that the European invasion has fixed polymorphism in large areas of the genome (16). This loss of genetic diversity is consistent with a strong bottleneck, but also with a selective sweep. Without independently rerunning the invasion of Europe from Asia, it is difficult to discriminate chance from selection.

Rust fungi are obligate biotrophic pathogens and this intimate association with a living host makes them ideal for the study of adaptation in agriculture (17). Rusts also make up some of the most devastating crop pathogens, infecting crops of global significance, such as wheat (18), soybean (19) and coffee (20). Rusts are known for their complex life cycles with multiple stages, in agriculture the main infection phase is clonal, where urediniospores are found erupting from pustules on the surface of leaves (Fig. 1A). Each spore contains two different haploid nuclei (dikaryotic). This phase can continue indefinitely on susceptible hosts but can also alternate with a seasonal sexual stage (17). While there are exceptions the general principle is that mutation increases polymorphism between dikaryotic content (and within the population) in the clonal phase and recombination shuffles beneficial variation in the sexual phase (17,18). The sexual phase of these pathogens may be a key determinant in their rate of adaptation. In most cases, the sexual phase of the life cycle occurs on a different host plant, termed heteroecious (as opposed to autoecious). In the case of wheat stem rust this alternate host is barberry which has been associated with increased rust virulence in Europe and the US for hundreds of years (21).

**Figure 1.**
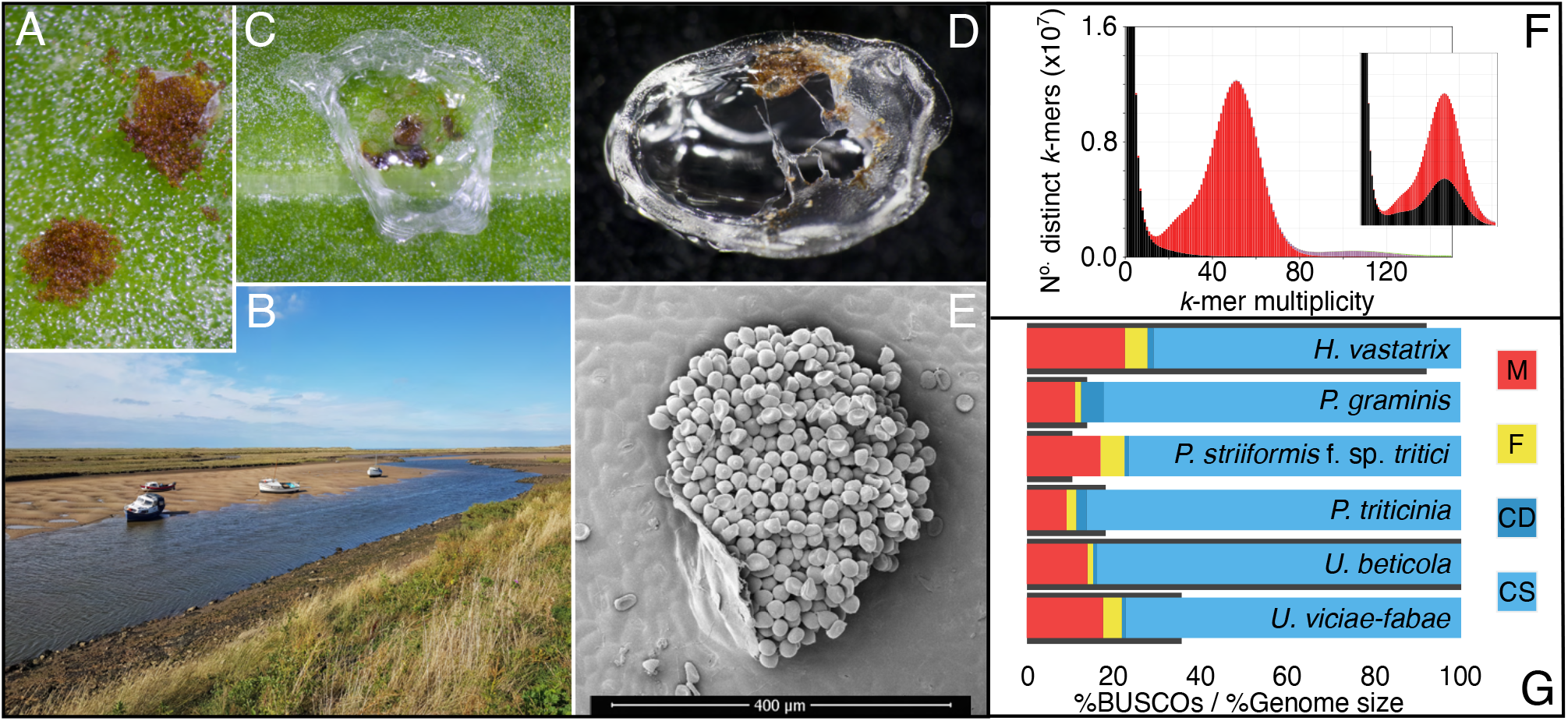
Beet rust, isolated from wild and agricultural hosts, peeled and sequenced. (A) *U. beticola* pustules (2-4mm approximately). (B) Sea beet found along an estuary, from which leaves are clipped and brought back into the lab. (C) Pustules are covered with 5μl peel solution which sets and is peeled off for library preparation and sequencing (D). (E) An electron micrograph image of a rust pustule (artificially coloured), individual urediniospores are visible. (F) Rust genome *k*-mer spectra is a histogram demonstrating the number of *k*-mers (in the reads) found at a given multiplicity (or depth). *K*-mers present in the reads but absent in the assembly are plotted in black, present once in the assembly in red, then purple for twice. Importantly, rare *k*-mers (suspected errors) are not found in the main distribution of the assembly which centres around the sequencing depth (^~^50x). Content falls largely within this 50x peak (homozygous) with a slight heterozygous shoulder (^~^25x). The inset plot shows the *k*-mer distribution where all contigs without a blast hit are removed. The black peak in the main distribution suggests that this is real *U. beticola* content and should remain in the assembly. (G) BUSCO completeness scores are highlighted for *Hemileia vastatrix* (541.2 Mbp), *Puccinia graminis* (81.5 Mbp), *P. striiformis* f. sp. *Tritici* (61.4 Mbp), *P. triticinia* (106.6 Mbp), *U. beticola* (588.0 Mbp) and Uromyces viciae-fabae (209.5 Mbp). Colours indicate Missing, Fragmented, Complete and Duplicated and Complete and Single copy BUSCOs. Genome sizes are represented as grey bars that boarder BUSCO scores and are a percentage relative to the largest, *U. beticola*. The *U. beticola* genome while large has comparatively middle to low levels of missing, fragmented and duplicated BUSCO content.

In wheat yellow (stripe) rust the centre of genetic diversity and recombination occur in Nepal, Pakistan and China and yet their clonally sustained populations expand to every continent except antarctica (22). So, despite the knowledge that centres of diversity and recombination are important, these processes are difficult to study either because the locations of wild hosts are difficult to identify and sample and/or these centres of diversity are linked to the centre of domestication of the crop and difficult regions to access. In the present study, we selected a system specifically to investigate wild-agricultural adaptation with a requirement for an obligate biotrophic pathogen that lives on a wild crop relative. Sea beet (*Beta vulgaris* subsp. *maritima*) is a (largely self-incompatible) host found permanently along the western coastlines of Europe (23,24; Fig 1B) and is the wild progenitor of sugar beet (*B. vulgaris*). Sugar beet is one of the most recently domesticated crop species (circa 200 years, 25), it is grown for sugar production throughout Europe, where yield is impacted by its autoecious rust pathogen, *Uromyces beticola* (26).

This system is being developed to understand the nature and strength of the selective constraints arising from the wild and agricultural environments. Natural selection is expected to operate to distinguish genetic variation between wild and agricultural pathogen populations and that this should be more pronounced in genes important for success in these environments. Therefore, in rusts living on wild and agricultural hosts we should expect to see (1) population differentiation between isolates from the wild and agriculture and (2) within the genome, signals of diversification should be greatest at genes important for niche adaptation. To investigate whether the impact of adaptation to the agricultural environment is present in the genome of a fungal pathogen, we first assembled and annotated the *U. beticola* genome and characterised population genetic diversity in isolates sampled from sea and sugar beets across England. To facilitate genetic studies, we develop a method of peeling *U. beticola* spores from the surface of a leaf, allowing us to re-sequence and analyse variation across 1.87 million SNPs among 42 rust isolates from 24 sugar beets and 18 sea beets. Wild sea beets were sampled along approximately 370 km of coastline as were sugar beets (inland as the crow flies, ^~^200 km). Despite sea and sugar beet fungal samples sometimes being 10 km apart, we find evidence for two populations, split between an exclusively wild set of isolates and a group of predominantly agricultural isolates. Importantly, we find a signal of adaptive evolution in genes that putatively interact with the host (effectors) as well as a signal of increased levels of genetic variation at these effectors.

## Results

### A genome assembly for population genetics of U. beticola

In order to test our hypothesis, that adaptation to agricultural crops is evident in comparisons of wild and agricultural plant pathogen populations, we assembled a large rust genome (588 Mbp) into 19,690 contigs with a genome N50 of 74 Kbp, the largest contig is 554 Kbp . Rust genomes are often problematic to assemble because of the dikaryotic (n+n) content of the uredospore life stage which is sampled (26,27). Divergent haplotypic content would be evident as a double peak in the *k*-mer distribution of the reads (e.g. see Fig. 1 in ref 19). We do not observe this signal characteristic of divergent haplotypic content and instead our spectra plots suggest we have a genome with relatively low heterozygosity, misassembles or frame shifts (28; Fig. 1F).

We wanted to assess completeness of our *U. beticola* assembly and annotation so we used core gene presence in comparison to a range of other rust assemblies of smaller and equivalent size (20,29,30; Fig. 1G). BUSCO genome completeness places *U. beticola* (85.1% of the complete BUSCOs; 84.0% single copy) within the level of other rust assemblies. In addition, the level of duplicated BUSCOs is low in comparison to other rust assemblies where divergent dikaryotic content may be more problematic (Fig. 1G).

The *U. beticola* genome annotation identified 9,148 genes (17,591 transcripts) with an average transcript length of 2,057.7 bp (6.4 exons per gene) at a mean coding sequence (CDS, spliced from transcript in the annotation) length of 1240.9 bp (see also Supporting Information 1 & 2). A large proportion of genes were annotated as Transposable Elements (TEs, 41.4%, taking the total up to 15,612) within this repeat rich genome in which combined low complexity and interspersed repeats represent 89.96% of the genome. Our reassessment of published rust genomes shows that this level of repeat content is consistent with such a large rust genome (Supporting Information 3). Signal peptide information was used to define the secretome and then 225 effectors were identified using EFFECTORP2.0.

### In the UK rust is differentiated into two populations

We sampled and sequenced 46 individuals and after quality control 42 were used in population analyses. To assess population level diversity and divergence, we identified 1.87 million SNPs across the 588 Mbp genome from 42 individuals sampled from either a wild sea beet (*n*=18) or an agricultural sugar beet (*n*=24; Fig. 2A). Genetic diversity was used at two levels, using gene CDS regions for plotting a Neighbour-Net network and also using all SNPs to examine population subdivision based on discriminant analysis of principal components (DAPC; 31). The Neighbour-Net network showed differentiation of an exclusively wild group of individuals from three sites in the south-east of England (Fig. 2B; Table 1). Isolates in this wild clade were sampled from sites spanning approximately 80km, yet isolates from agricultural beets, sampled just 10km away, are found in the predominantly agricultural clade (Table 1, Orford). Despite these wild and agricultural isolates being found in close proximity this predominantly agricultural population extends across all agricultural sites, and also encompasses the two northern most wild sites (Fig 2A). Using all 1.87 million SNPs we used DAPC to assess population subdivision. All 41 PCs were retained initially to determine that there were two clusters or populations (*find.clusters*: 2-20) and then 15 PCs, accounting for 68.7% of the conserved variance, were used in the DAPC analysis (Fig. 2C). Individuals were sorted into two clusters which represent the southeast wild only group (*n*=13) and the majority agricultural group as identified by the network analysis of gene coding regions (*n*=29; see Fig. 2B,Table 1). The overall level of genetic differentiation (*F*_*ST*_) between these two populations is 0.113 (Fig. 2C) and given the proximity (and spread) of wild and agricultural populations the processes driving this divergence may be reproductive or based on natural selection.

**Table 1.**
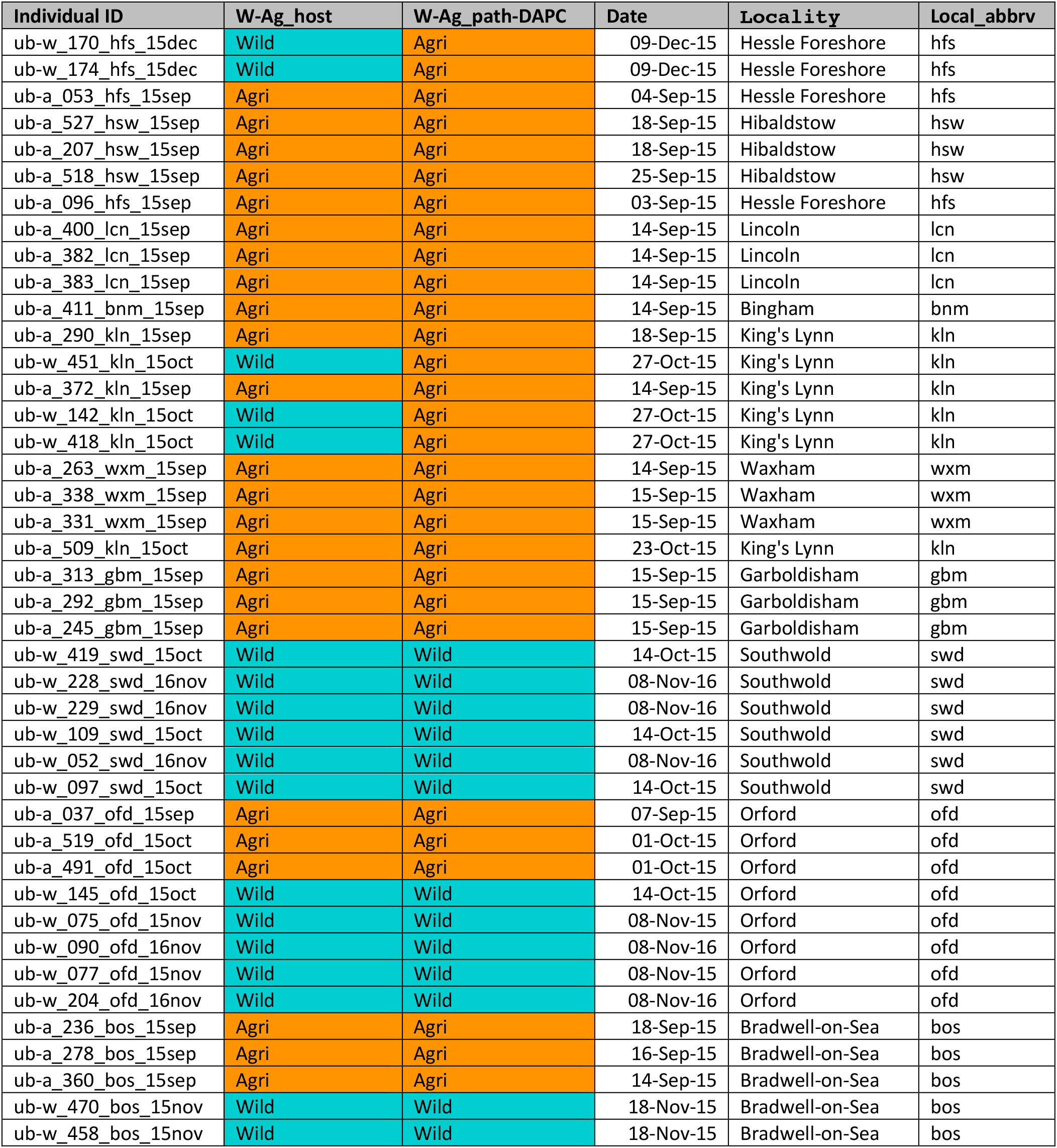
Isolate sampling data listed north to south including information on whether isolation was from a wild or agricultural beet (W-Ag_host) and how isolates clustered after DAPC analysis of genotypes (W-Ag_path-DAPC). Local abbreviation and colour (W-Ag_host) is used in samples names in Figure 1.

**Figure 2.**
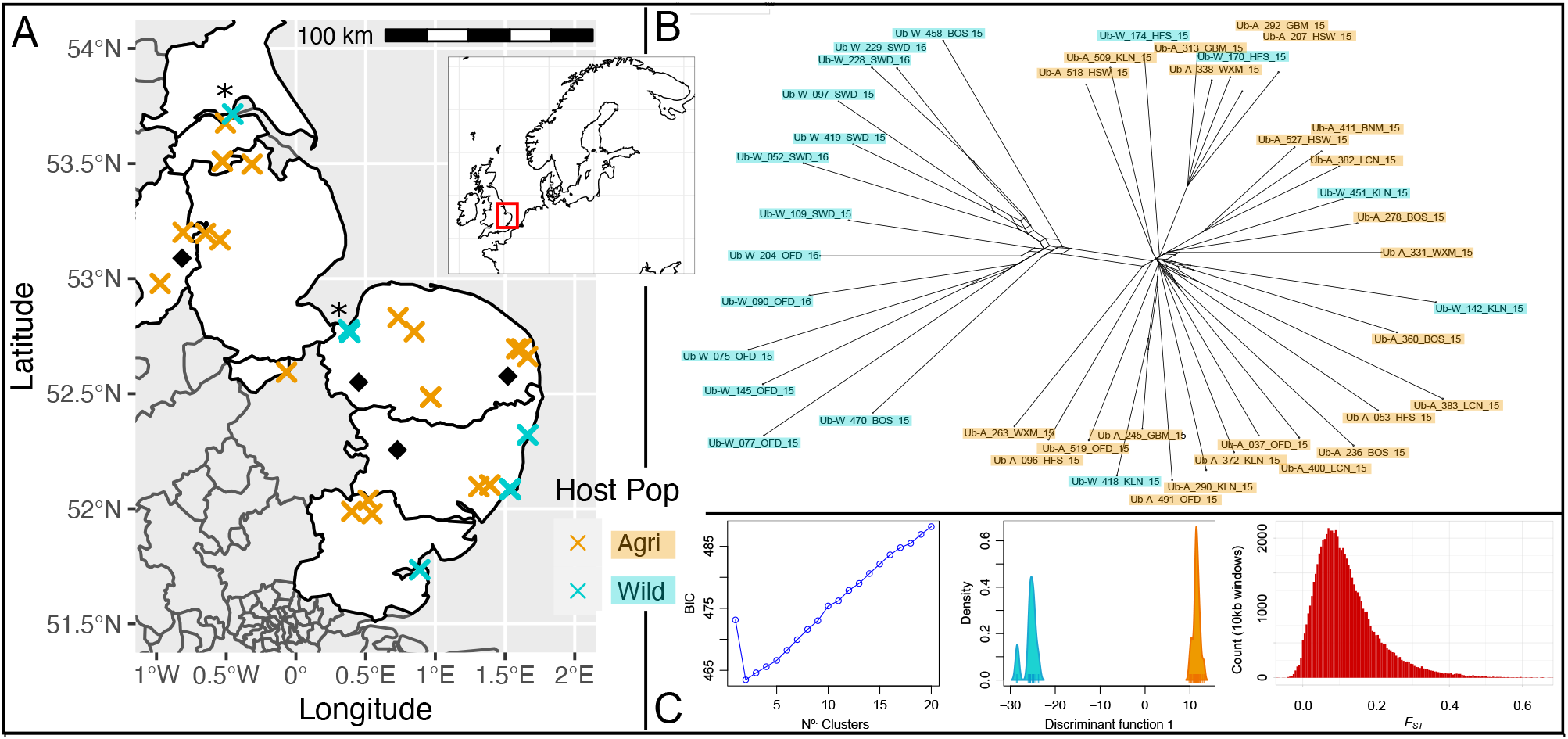
Population differentiation separates an entirely wild population from all agricultural and remaining wild individuals. (A) Map of the UK shows samples collected from wild (blue crosses) and agricultural (orange crosses) hosts. Overlapping crosses hide wild samples which can be distinguished using Table 1. Norther wild samples belong to the Agricultural group and are accompanies by an asterisk. Diamonds indicate beet factories. (B) SplitsTree network generated using the CDS regions of 15,473 genes (15.2 Mbp) shows a clear differentiation of the wild only population. Heterozygous sites coded using IUPAC ambiguities. From here onwards when referring to populations we refer to the grouping observed in this network & by DAPC. (C, left to right) The optimal number of clusters was two based on the lowest associated BIC and, given just two clusters, the blue-orange distribution shows how individuals cluster across the single discriminant function. The genome wide genetic differentiation between clusters, *F*_*ST*_=0.113. However, the red distribution shows genetic differentiation in 10 Kbp windows across the genome and emphasises that differentiation can vary. Selection against polymorphism moving from the wild to agriculture (or *vice versa*) would drive genes towards an increasing *F*_*ST*_.

### Reproduction may be partitioned differently in wild and agricultural populations

In order to understand whether the relative levels of sexual and asexual reproduction are different between wild and agricultural populations we assessed the level of inbreeding. The inbreeding coefficient, *F*_*IS*_, describes the proportion of genetic variation contained within an individual relative to its subpopulation. The measure of *F*_*IS*_ most often scales between zero and one and indicates random mating and inbreeding, respectively. In the present data, while we do not observe a significant difference in the level of heterozygosity per individual, we see that *F*_*IS*_ is negative in all agricultural individuals (Fig. 3A & 3B). A negative value of *F*_*IS*_ suggests excess heterozygosity and a role for the preservation of polymorphism via some clonal reproduction. Although the wild population also contains individuals with negative *F*_*IS*_ values, in general, the wild population have higher *F*_*IS*_ values that are closer to and above zero (Fig. 3B). The wide distribution of *F*_*IS*_ values in the wild suggests the occurrence of both clonal and sexual reproduction.

**Figure 3.**
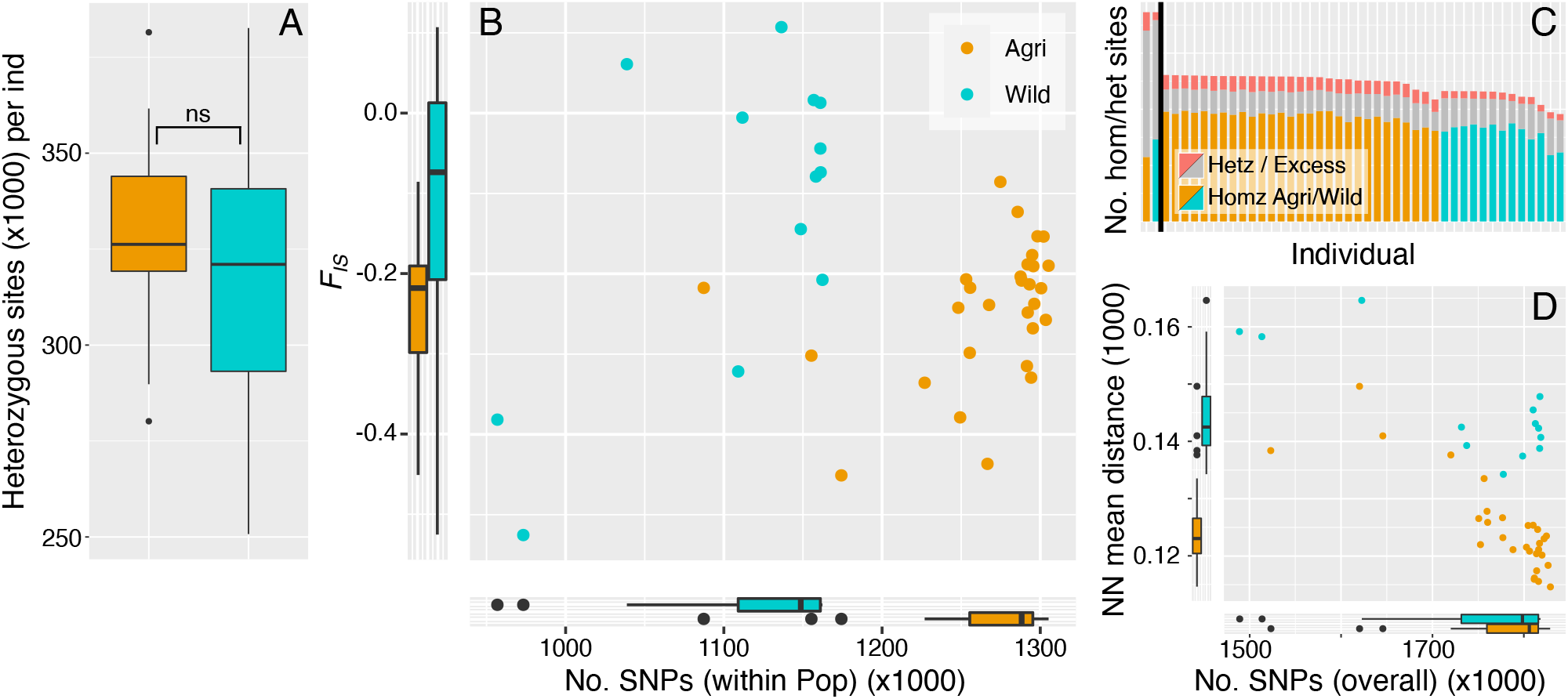
Individual level polymorphism is different between agricultural and wild populations. (A) There is no significant difference between the number of heterozygous sites in agricultural and wild individuals (median heterozygous site No.: Agri=326,246, Wild= 321,031; Wilcoxon W=225, p=0.332) and (B) wild rust isolates have fewer SNPs (within population) and broadly distributed *F*_*IS*_ values that tend more towards zero and above on average. Agricultural isolates have larger numbers of SNPs and *F*_*IS*_ values that are more negative on average. There is a significant difference in the *F*_*IS*_ values of agricultural and wild populations (Median *F*_*IS*_: Agri=−0.21802, Wild=−0.07371; Wilcoxon W=92, p-value=0.008). Negative *F*_*IS*_ values are indicative of an excess in heterozygous sites and consistent with clonal modes of reproduction, as are accumulation of mutations at the individual level. (C) Numbers of homozygous and heterozygous SNPs per individual. Individuals represented by a single bar where homozygous sites coloured by population (orange agri and blue wild). Grey represents the number of heterozygous sites per individual with red tips indicating the proportion of those heterozygous sites that are in regions of excess heterozygosity. The vertical black line separates population total values (left, reaching 1.87 million SNPs) from individual values (right). (D) Individuals observed with fewer SNPs within population (compare B x-axis) are not less divergent overall, plotted against mean Neighbour-Net distance (see Fig. 2B).

In order to specifically investigate whether the heterozygosity associated with the inbreeding coefficient could have been caused by other processes we looked for an association with other features of the genome. As well as clonal reproduction, excess heterozygosity can also be caused by processes such as genome organisation and repeat content driving erroneous heterozygosity via read miss-mapping. Blocks of excess heterozygosity (at the 5% significance level based on Hardy-Weinberg Equilibrium test) impact agricultural and wild populations to different extents. We find that the agricultural population has approximately 22 thousand blocks (two or more excess heterozygosity sites) where the wild has just less than ten thousand blocks (mean block length: Agri=227.7 bp, wild=124.0 bp; mean No. excess heterozygous sites: agri=5.1, wild=5.9). While the agricultural population has been impacted by excess heterozygosity to a greater extent, we found no association of these regions of excess heterozygosity with repeat or genic regions and so it is difficult to discriminate between effects of genome structure from mode of reproduction.

### Effector genes provide evidence for adaptation to agriculture

We have identified a wild-only population of isolates as distinct from another population that infects all agricultural hosts sampled, plus five northern wild hosts. Despite the presence of some wild infecting isolates in our ‘*agricultural’* group, potentially dampening these effects, we set out to identify signals of adaptive variation in CDS regions and divergence in effector genes between these wild and *agricultural* groups.

Natural selection operates on variation already present in a population and so here we measure levels of nucleotide diversity (*π*). Our first observation is that a combination of short CDS regions and the level of polymorphism in genes results in less than half of all gene CDS regions being polymorphic (43.4%; Fig. 4A). At the genome scale (10 Kbp windows), levels of nucleotide diversity in the wild population are marginally (but significantly) higher than those of the agricultural population (Fig. 4B). For those effectors and non-effector genes that contain polymorphism within their CDS region, we observe first, that genetic diversity in non-effector genes is higher in the wild than in agriculture, second, that effectors are more polymorphic than non-effectors in both populations and third, that effector diversity in the wild is greater than that present in agriculture (Fig. 4C; Supporting Information 4).

**Figure 4.**
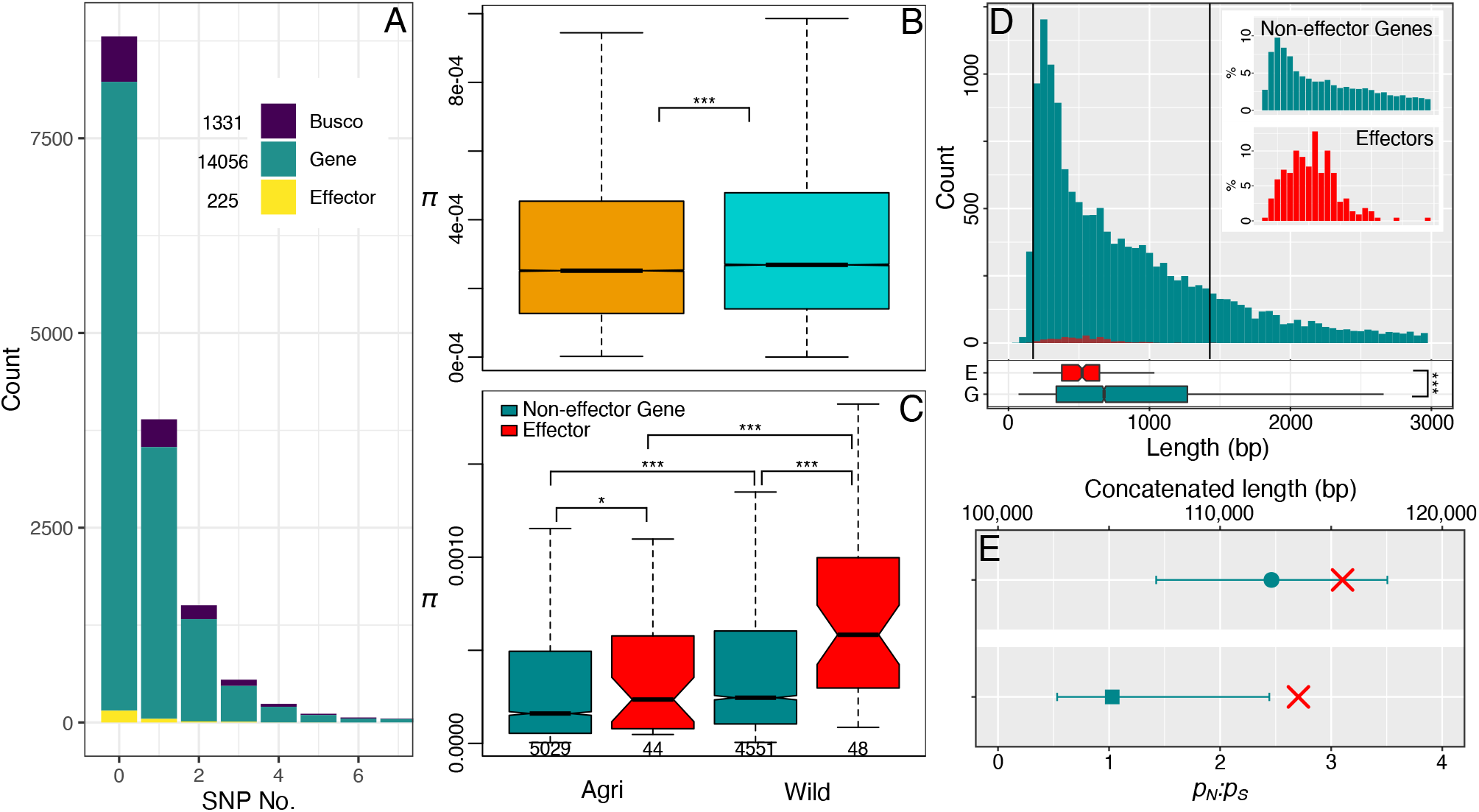
Wild effector genes maintain a signal of adaptive diversity. (A) Histogram of genes with a given number of SNPs in their CDS region (between 0-7 SNPs). 8808 genes have zero SNPs within their CDS region and 6744 have one or more SNPs. (B & C) Genome wide levels of nucleotide variation (*π*) are higher in the wild papulation (B, median *π* 10 Kbp windows: Agri=0.251×10^−3^, Wild=0.269×10^−3^; Wilcoxon(*π*) W=2440369930, p<0.001). Considering polymorphic genes, this pattern of wild maintenance of diversity is also observed where, Agricultural non-effector genes have significantly less diversity than Wild non-effector genes (C, median *π* non-effector genes: Agri=0.161×10^−3^, Wild=0.245×10^−3^; Wilcoxon(*π*) W=9454106, p<0.001). Despite low numbers of effectors with polymorphism in the CDS (see gene *n* under boxplot in C) maintenance of polymorphism in these host interaction genes is significantly higher than that of non-effectors within both agricultural and wild populations (C, median *π* effectors: Agri=0.236×10^−3^, Wilcoxon(*π*) W=93488, p=0.038; Wild=0.583×10^−3^, Wilcoxon(*π*) W=69958, p<0.001). Finally, wild effectors are significantly more polymorphic than agricultural effectors (Wilcoxon(*π*) W= 638, p<0.001). (D) Histogram of observed effector and non-effector gene CDS lengths (between 1-3000 bp) shows counts and percentages (insert) per gene type. Vertical black lines indicate the effector size range, and the insert shows the distribution of sizes within that range (median CDS length: non-effector gene= 678, effector= 522; Wilcoxon(bp) W=2094583, p<0.001). (E) Point and error bars show the Length and *pN:pS* values after concatenating effectors (observed, in red), or size matched non-effector genes (bootstrapped, in turquoise). Size matched non-effector genes are no longer than effectors (effector Length=116 Kbp; non-effector gene median (5-95%CI) Length=112 Kbp (107 Kbp – 118 Kbp); randomisation test p=0.148). However, size matched non-effector genes have significantly lower levels of adaptive diversity than effectors (effector *p*_*N*_:*p*_*S*_=2.71; non-effector gene median (5-95%CI) *p*_*N*_:*p*_*S*_=1.029 (0.533 - 2.443); randomisation test p=0.043).

Next we used a measure of the average number of non-synonymous to synonymous polymorphism (*p*_*N*_:*p*_*S*_) within non-effector genes and effectors as a whole. However, as mentioned above, many CDS regions contain zero SNPs and many others contain just a single SNP. A single SNP makes calculating adaptive diversity difficult in cases where genes have a single non-synonymous polymorphism (*p*_*N*_), as the ratio of non-synonymous polymorphisms (to synonymous, *PS*) is infinity. This process has the potential to impact effectors to a greater extent as these tend to be shorter on average than non-effector genes (Fig. 4D). In order to account for this, we used a concatenation and bootstrapping approach to compare adaptive diversity between effector and non-effector genes. This approach involved concatenating all 225 effectors to measure a single *p*_*N*_:*p*_*S*_ value (averaged pairwise among alleles). Not only does this solve the problem of single *P*_*N*_ polymorphisms but it also utilises regions of zero polymorphism in other genes CDS regions. This process was then repeated (×1000 with replacement) for size matched non-effector genes. This demonstrated that the signal of adaptive diversity was greater in Effectors than non-effector genes of the same size (Fig. 4E).

After demonstrating that diversity is greater in the wild and that effectors carry an increased signal of adaptive diversity we next wanted to test whether effectors were more different between wild and agricultural populations than other genes. We predicted that host genetic diversity is one of the largest discriminating factors between the wild and agricultural environment and therefore we expect to observe it in the genes that putatively interact with the host, the effectors. Here we measured the genetic differentiation at all genes which included 5 Kbp flanking each gene and found indeed that effectors were significantly more diverged than non-effector genes (Fig. 5).

**Figure 5.**
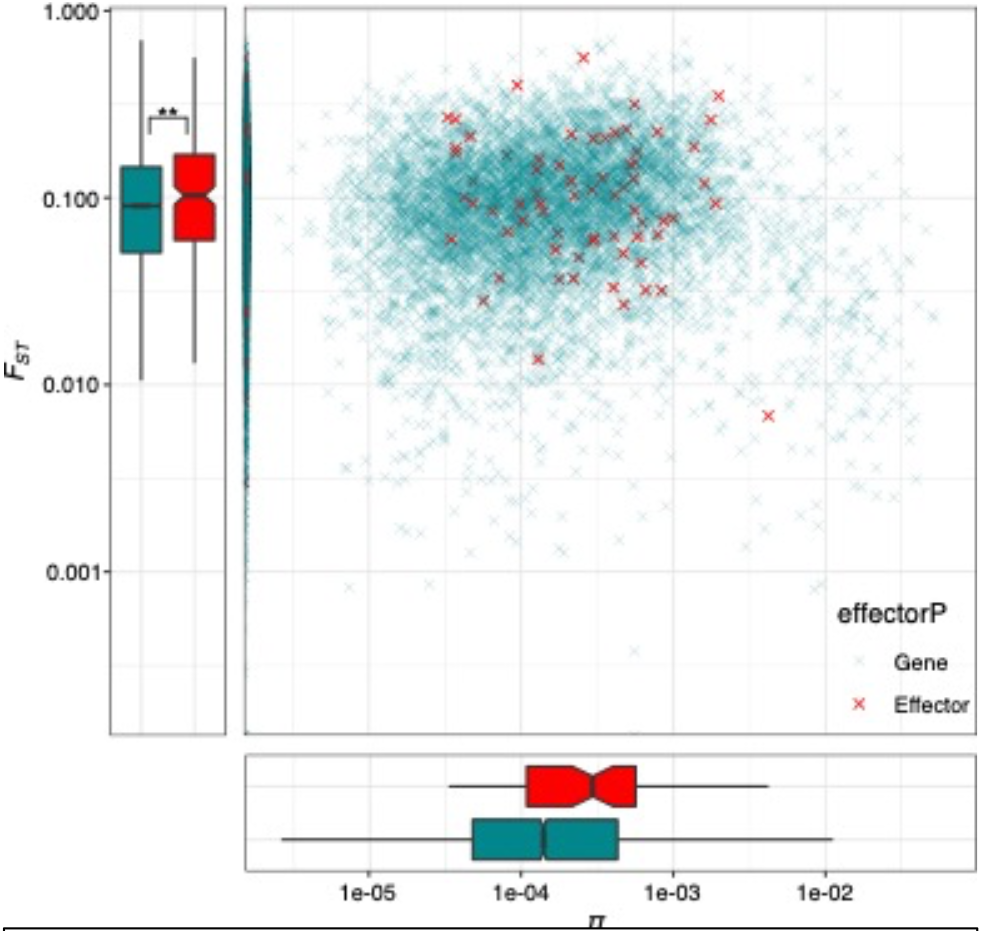
Evidence for adaptation to agriculture (and the wild) is present in effector diversity. Levels of genetic differentiation between the agricultural and wild populations non-effector genes and effectors. effectors are significantly more differentiated between populations than non-effector genes (median *F*_*ST*_: non-effector genes= 0.089, effectors= 0.100, Wilcoxon(*π*) W= 1515702, p=0.005).

## Discussion

### A genome for population genetics using peel sequencing

Based on our wild-agricultural adaptation hypothesis we predicted that we would observe genetic differentiation between wild and agricultural populations. We set out to establish this using peel-sequencing, a method developed to allow us to paint on and peel off (and genome sequence) rust pustules from the surface of a leaf. Our peel method reduces the level of sequencing resource devoted to the host and our preliminary results were generated prior to sequencing our draft genome and instead, we assembled and mapped to a peel genome. This preliminary genome was approximately one third the size of our draft but it allowed us to estimate the genome size and sequencing effort required for resequencing as well as highlighting the main signal of our analyses (without a draft reference genome; data not shown). However, we subsequently generated a reference assembly and annotation using DNA and RNA from *U. beticola* spores combined from multiple pustules on a leaf. The *U. beticola* genome at ^~^600 Mbp, is large for a fungus but not for a rust (19). The rusts have dikaryotic haploid nuclei and in many rusts this produces a scenario (Meselson effect) in which extended periods of clonal reproduction (without recombination) increase the divergence of homologous content (18,32). However, many agronomically important rusts also have an alternate host which is used by the fungus to enter its sexual phase (27). In the case of the wheat rusts, hundreds of years before a causative link was established, superstition drove the removal of its alternative host, *Berberis*, from cereal growing regions (21). *U. beticola* reproduces on wild sea beet and, the permanent proximate availability this host offers a (*post hoc*) explanation as to why we have not observed divergence of karyon (reduced heterozygosity) in *U. beticola*.

Agriculture provides the largest annually replicated plant pathogen experiment on Earth. Replicated genotypes are put out annually and pathogens must invade from a reservoir and adapt. Natural selection acts upon genetic variation that is already present in a population and the idea that pathogens evolve in reservoirs and then reinvade new populations or species is neither new nor limited to plants and fungi, viral zoonoses are particularly pertinent to human health (e.g. HIV, SARS-CoV2, 33). Farming is a relatively recent phenomenon (by humans, 34) which means that plant pathogens must either still be sharing genetic variation with pathogens on progenitor hosts or, they are reproductively isolated from them and specialising. Either way, we can use the signals of differentiation and divergence to understand the genes important for adaptation to agriculture and perhaps more importantly, the processes underlying adaptation. Here we investigate these processes by developing the wild-agricultural beet rust system (*Beta vulgaris* subsp. *maritima*, *B. vulgaris* and *Uromyces beticola*, respectively). Consistent with predictions that come from our hypothesis on how wild-agricultural pathogens evolve, we show that the signal of adaptation is found to a larger degree in fungal genes that putatively interact with the host.

By assembling a draft genome and developing a peel sequencing method that allows preferential sequencing of pathogen to host material, we first identify two genetic clusters of rust in England, a population found solely on wild host plants as well as a population found predominantly on agricultural hosts. Levels of genetic diversity are marginally higher in the wild population although agricultural individuals tend to carry greater individual level polymorphism. Rust pathogens can reproduce clonally or sexually and despite not identifying any direct clones in our clone correction analysis we observed a signal of increased levels of heterozygote excess. This finding is consistent with an increase in clonality in agriculture. Next, we established that genes predicted to interact with the host, effectors, carry an increased level of amino acid changing polymorphisms compared to non-effector genes. Finally, we identify that on average these effector genes are more differentiated between wild and agricultural populations than non-effector genes. This finding is important for all crop pathogen systems in identification of genes important for the adaptation of pathogens to agriculture but more generally, also provides a framework for identification of agriculturally adapted genes as well as a means to go on to identify the mechanisms that facilitate that adaptation.

### Genetic divergence between agricultural* and wild populations

To assess the genetic differentiation between wild and agricultural isolates, we chose a mix of isolates sampled from wild and agricultural hosts. Wild samples were spread latitudinally along the regions from which we received agricultural samples from growers. Discriminant Analysis of Principal Components highlighted two distinct clusters of isolates in England, a group that lives exclusively on wild hosts and a group that lives predominantly on agricultural hosts (24 out of 29 isolates). The wild population was found at three sites in the east of England (distributed across ^~^80 km). The isolates identified within the agricultural clade spanned the wild populations and came within 10km at the nearest wild sampling site. This predominantly agricultural clade was identified for all isolates found in agriculture, plus five isolates found on wild hosts in two more northern sites. We can’t yet account for these wild infecting *agricultural* isolates and we go on to analyse the genetic diversity and divergence of these two populations (wild and agricultural compatible) with an aim to address the role of host genetic diversity and broader metapopulation dynamics in our observation.

Investment in clonal or sexual reproduction can vary with environmental heterogeneity, where the advantages of sex for rapid adaptation diminish to fecundity as environmental heterogeneity declines (35,36). This drives the prediction that agricultural pathogen populations may increase the relative rate of clonal to sexual reproduction to rapidly infect hosts before cropping (or chemical treatment) and not pay the cost of reduced rate of adaptation because hosts are genetically more similar than wild hosts (22). Indeed, we do observe, using an inbreeding statistic (*F*_*IS*_), that the agricultural population has an increased signal of clonality (37). This is a consequence of excess heterozygosity which is more prevalent across the agricultural genome. In wheat stripe rust, bouts of clonality spanning decades were used to account for increased levels of heterozygosity and is believed to be driven by the lack of a sexual host (18). We see excess heterozygosity not increased heterozygosity, although our power to detect increased heterozygosity in the agricultural population may be impacted by the low numbers of individuals assigned to our wild clade.

Excess heterozygosity could also have been caused by assembly and mapping errors, particularly in this repeat rich genome. The agricultural population has both a greater number of regions impacted by excess heterozygosity as well as a larger overall proportion of the genome found to be in heterozygote excess. However, blocks of heterozygosity appear no more associated with repeat regions as they do with genic regions. It is worth adding that the reference genome was generated from an agricultural isolate and this perhaps reduces the chances of heterozygosity caused by repeat differences and mapping errors. A long-read assembly, spanning repeat regions, would allow us to convincingly address the impact of repeats and delineate, differential agricultural genome structure, from clonal reproduction in the agricultural population.

### Effector genes carry a signal of adaptation to Agriculture

Effector proteins must both avoid detection by the host immune system as well as target host gene products (which are also evolving) and so they must continually adapt in order to avoid detection and remain effective (38,39). In the present study we categorised those gene products that are secreted outside of the fungal cell and used machine learning to classify 225 genes with effector properties. We use those genes as a group with the expectation that their evolution will allow us test our hypothesis: that natural selection should operate to further distinguish genes important for success among wild and agricultural environments.

We set out to first, establish the presence of signals of adaptation in effectors as compared to non-effector genes. Consistent with predictions of linkage to loci operating under balancing selection (40), nucleotide diversity is significantly higher in effectors than in non-effector genes of the same population. Moreover, the diversity of polymorphic wild effectors is twice that of other gene categories or the genome average. These diversity analyses are all done on CDS regions and it is worth pointing out here that of the 225 effectors identified only 44 and 48 (Agri and Wild, respectively) contain polymorphism in this region. Despite these reduced numbers of polymorphic CDS regions we retain enough power to show that differences between effectors and non-effector genes are statistically significant. It is because of these low numbers of polymorphic CDS regions that for analysis of adaptive diversity, which can only use synonymous and non-synonymous polymorphism (*p*_*N*_:*p*_*S*_ in the CDS), we compared observed mean *p*_*N*_:*p*_*S*_ of all effectors concatenated, to bootstrapped (size matched) non-effector genes. We found that effectors contained a greater level of adaptive diversity than expected based on that of non-effector genes. Therefore, in addition to diversity in general, adaptive diversity is higher in these host interaction genes.

Finally, with effector genes carrying a signal consistent with their evolution and adaptation to host resistance, we test for a signal of specificity to the wild and agricultural environments and indeed we find that genetic differentiation in effectors is significantly higher than that of non-effector genes. Again, from the proximate sugar beet rust invasion perspective, those effectors that are more highly divergent between wild and agricultural populations represent those *agriculturally adapted* variants whose mode of action is most important to elucidate. However, more broadly these analyses provide a framework by which we may identify genes that are adaptively divergent between the wild and agricultural populations and are therefore, important agricultural adaptation genes.

In the present study we identified the presence of signals consistent with agricultural pathogen adaptation to agricultural hosts. This finding has implications in the beet rust system but, more importantly also for the way we understand and analyse pathogen evolution. We intend to use this system to further explore wild-agricultural plant pathogen evolution. This first look at population genetic diversity raises questions on the role of broader metapopulation dynamics as well as whether this signal is replicated across other wild-agricultural boundaries, in Germany and France for example. The present work also raises questions about the role of linkage, repetitive elements and genome organisation in local adaptation. The rusts are a problematic lineage of crop pathogens with intricate and complicated haploid, diploid and dikaryotic life stages (27,41). Many other fungal and insect systems partition their reproduction and favour alternative hosts for sexual reproduction and this strategy may be particularly advantageous in a system where the crop host is not present year-round. Clonal reproduction suits a boom-and-bust invasion lifestyle, but a bit of sex goes a long way, and it is perhaps this ability to partition these modes of reproduction on different hosts that has made them so adept at capitalising the crop system. Ultimately, this study offers a beginning to understand multi -host -pathogen systems that might one day help predict sources of adaptation in reservoir systems and inform targeted treatment of agriculture to reduce adaptive introgression and increase the durability of the resistance already in the system.

### Materials & Methods

In order to test our hypothesis, that pathogen adaptation to agricultural crops will be evident in comparisons of wild and agricultural plant pathogen populations, we identified a plant pathogen, a rust fungus (Uromyces beticola) that is an obligate biotroph found in the UK living on both wild sea beet and agricultural sugar beet. We sampled isolates in both the wild and agriculture, sequenced and assembled a large (agricultural) fungal genome, sequenced expressed genes for genome annotation and then re-sequenced 46 wild and agricultural rust isolates for population genetic analysis, of which 42 made it through quality control.

### Genome Assembly and annotation of U. beticola

A rust infected sugar beet plant from Norfolk was brought into the lab to allow the infection to progress in the absence of agitation by wind and rain. Multiple pustules from a heavily infected single leaf (Fig. 1) were broken and spores collected in order to extract DNA (see below for CTAB details) for DISCOVAR PCR free library preparation and sequencing (42). A HiSeq2500 was used to sequence 43.8 Gbp of data which was estimated to be 73x coverage based on genome size estimates of ^~^600 Mbp from earlier tests of peel sequencing. Post assembly, contigs less than 1 Kbp were removed as they don’t represent a real-terms increase in the span of a single read pair. We used ABYSSv2.0.2 (43), KATv2.3.4 (28), BLOBTOOLS v0.9.19 (44) and BUSCOV4.0 (45; against basidiomycota_odb10) to assess genome content, contiguity and completeness. BLOBTOOLS was used to retain contigs with BLAST hits to Basidiomycetes as well as those without a hit.

To generate genome annotation, RNA was extracted from rust infected sugar beet leaves collected in the field. Using a soft bristled toothbrush, pustules were lifted/brushed from each leaf into a 2ml microfuge tube with the aim of combining 100mg of material (×10 tubes). Spores were then flash frozen in liquid nitrogen in preparation for RNA extraction (10 samples). In addition, leaf punches were also taken; punches were positioned at pustules away from vascular elements of the leaf in order to capture fungal expression *in planta* (2 samples). RNA was extracted using the Qiagen AllPrep Fungal DNA/RNA/Protein Kit and stranded libraries were prepared using the NEBNext Ultra II Directional with poly-A selection. Libraries were sequenced on one lane of the NovaSeq 6000, SP flow cell with the 300 cycle kit (150 bp PE) with v1 chemistry and reads were quality controlled using CENTRIFUGE (46).

Gene models were annotated using a workflow which incorporated repeat identification, RNA-Seq mapping and assembly, and alignment of protein sequences from related species (Supporting Information 1). Alternative reference guided assembly methods were employed (Scallop, 47, StringTie2, 48) to assemble transcripts for each sample. From these a filtered set of non-redundant transcripts were derived using Mikado (49). Gene models were classified based on alignment to protein sequences, identifying the subset of gene models with likely full-length ORFs. The classified models together with aligned proteins and repeat annotation are provided as hints to AUGUSTUS (50). Three alternative AUGUSTUS gene builds were generated using different evidence inputs or weightings. These together with the gene models derived from the Mikado transcript selection stage were consolidated into a single set of gene models using Minos (49). The Minos pipeline scores alternative models based on the level of supporting evidence (protein homology, transcriptome data) and gene structure characteristics (e.g. CDS, UTR features) to select a representative gene model and alternative splice variants.

In plant pathogenic fungi, gene products that are secreted outside of the fungal cell and into the host are considered candidate host interaction genes, potentially facilitating infection. These putative effectors are identified here using signal peptide information, genes with the presence of a signal peptide using SIGNALP v3.0: (-s notm -u 0.34; ,51) and the absence of transmembrane or mitochondrial localization signals using TMHMM v2.0 and TARGETP-2.0 Server (52) were finally assessed for sequence similarity to known effectors using EFFECTORP2.0 (53). Assessment of putative effector (henceforth, effector) density was done using the method of Raffaele *et al.* (54).

### Uromyces beticola isolate collection and sequencing

Samples were collected over the period of September to December in 2015 and 2016. Wild sea beet rust samples were collected from UK east coast sites between Southminster and Hull (^~^370km of coastline) and agricultural sugar beet rust samples were posted to the Earlham Institute (via the British Beet Research Organisation, BBRO) by beet growers covering approximately the same latitudes (^~^200km). DNA extraction from single rust pustules was carried out at the Earlham Institute (EI) using the peel extraction method. This method involves pipetting 5μl cellulose acetate onto a rust pustule, allowing one or two minutes to air dry and then peeling the fungus away from the leaf (Fig. 1C,D) and depositing the peel into a tube for storage (−80°C) prior to agitation and modified phenol-chloroform DNA extraction. This method maximises pathogen to host DNA in the extraction and avoids using excess sequencing resource on the host genome.

Peels were ground using one 4 mm and five 1mm stainless steel ball bearings in a TissueLyser (Qiagen, Valencia, CA) for 60 seconds at a frequency of 22 Hz. Fragmented peels were then incubated at 50°C for 1 hour in 500μl extraction buffer (2% cetyltrimethylammonium bromide, 1.4 M NaCl, 20 mM EDTA (pH 8), 100 mM Tris-HCl (pH 8), 0.2% β-mercaptoethanol, 1 mgml^−1^ proteinase K (55). Phenol:Chloroform:Isoamyl Alcohol 25:24:1, saturated with 10mM Tris, pH 8.0, 1mM EDTA (500 μl) was added to each sample and vortexed before centrifugation at 14,000g for 5 min. The aqueous (upper) phase (^~^450 μl) was harvested into a fresh microcentrifuge tube. To this 100% (v/v) of Agencourt AMPure XP (Beckman Coulter™) or a homemade mix Sera-Mag Speed-beads (Fisher Scientific, cat. #65152105050250) in a PEG/NaCl buffer beads (Rohland and Reich 2012) magnetic beads was added, samples were vortexed for 20s and incubated for 10 minutes at room temperature. The tubes were placed on a magnetic stand, and supernatant removed and discarded after all of the beads had been drawn to the magnet (^~^2 min). The beads were washed three times with 1 ml of 80% EtOH. After the last wash, the EtOH was removed and the beads allowed to air dry for 5 minutes. Magnetic beads were then resuspended in 55μl of EB buffer (10 mM Tris-HCl) and tubes incubated at 37°C for 10 minutes (vortexing every 2 min). To the eluate 1 μl of 10% (v/v) RNase A (100 mgml^−1^) was added before incubating at 37°C for 30 minutes. Magnetic beads (50μl) were added to each sample, vortexed and incubated at room temperature for 10 minutes. Sample tubes were then placed onto a magnetic rack for 2 minutes and the supernatant discarded. Beads were washed twice with 200μl of 80% EtOH. Ethanol was removed and the beads left to air dry for 5 min before resuspending in 55 μl of TLE buffer (Tris low EDTA – 10 mM Tris-HCl, 0.1 mM EDTA). The tubes were incubated at 37°C for 10 minutes (vortexing every 2 min). The tubes were again placed onto a magnetic rack for 2 min and the cleared eluate was harvested and stored at −20°C.

Libraries were prepared using the LITE method (56) and 46 isolates were genome sequenced at the Earlham Institute on nine lanes of an Illumina HiSeq4000 (150PE) generating ^~^750 Gbp of read data and an estimated mean depth of 27.5x per individual.

### Mapping and SNP calling

Reads were quality trimmed (minimum length 90, quality 30, --paired; TRIM_GALORE v0.4.0; Babraham Institute, Cambridgeshire, UK). BWA mem (57) was used to map reads and SAMTOOLS v1.5 (58) and BCFTOOLS v1.3.1 were used to sort and remove duplicate reads and mpileup (-t DP) to call variants (bcftools call -c). VCFTOOLS v0.1.13 (59) was used to filter SNPs to an individual minimum depth of five, maximum depth of 1.8 × mean depth per individual and a minimum genotype quality of 30. SNP sites with more than two alleles were excluded as probable errors. Finally, sites that were missing in 30% or more individuals were also removed. Four individuals were removed from further analysis because their average depth was less than 10x leaving 42 individuals for population analyses.

### SNP diversity and divergence analysis

Population differentiation was determined using Discriminant Analysis of Principal Components (DAPC, Adegenet; ,31,vcfR v1.12.0; ,60) based on all 1.87 million SNPs. Initially, the *find.clusters* function was used with all 41 principal components in order to identify two genetic clusters based on the lowest Bayesian information criterion (BIC). DAPC was then run using 15 principal components accounting for 68.7% of the conserved variance. Clone correction (*mlg.filter*) was conducted using a filtered SNP dataset using 1 SNP per 100 Kbp (1.1% of the total) and no clonal genotypes were condensed in the dataset regardless of the threshold used from *filter_stats* (farthest, average or nearest). All 42 individuals were retained for further analysis and plotted on the map of the UK using rnaturalearth.

VCFTOOLS was used for window analyses (10 Kbp windows), *F*_*ST*_ analyses of gene regions was based 5 Kbp up- and down-stream of gene boundaries. DNASP v6.12.01 (61) was used to calculate population genetic statistics per CDS after two levels of phasing, first using phase informative reads and then using linked homozygous SNPs to infer haplotypes from population data (SHAPEIT v2.20: assemble --states 1000 --burn 60 --prune 60 --main 300 --effective-size 88000 --window 0.5, 62). The effective population size estimate (*θ*=*4Neμ*) for SHAPEIT was calculated using mean *π* from 50kb windows from across the genome (*π* = 0.000352393) using an assumed mutation rate, *μ* = 1×10^−9^ (63). Ten contigs (19 genes) failed the phasing step and were removed from further analysis. Fasta conversion from vcf (GATK v3.5.0 - FastaAlternateReferenceMaker, 64) was done for spliced CDS regions (Cufflinks v2.2.1 -gffread, 65) per individual for all genes with read coverage greater than 60%. Analyses and plotting were done in R v3.6.3.

YN00 (PAML v4.9, 66) was used to calculate the average pairwise *p*_*N*_:*p*_*S*_ ratio per gene (nonsynonymous polymorphism, *p*_*N*_ and synonymous, *p*_*S*_). A gene concatenation and bootstrapping procedure was also used in order to recover a measure for genes with low diversity. The *p*_*N*_:*p*_*S*_ ratio can’t operate on genes with a single non-synonymous mutation and this scenario tends to impact short genes. To address this, we concatenated all effectors per individual and measured a single mean observed effector *p*_*N*_:*p*_*S*_. We then ran a bootstrap sampling regime in which we first, sampled effectors (with replacement) and binned them into length categories of 50 bp intervals. Second, we sampled from the non-effector gene set (with replacement) into the effector size bin frequency distribution. We concatenated those genes, measured their length and mean the *p*_*N*_:*p*_*S*_ ratio. This procedure was replicated this 1000 times in order to generate null distributions for concatenated *p*_*N*_:*p*_*S*_ ratio and length for non-effector genes.

CDS gene diversity data for 15,612 genes from DNASP & PAML are combined into a single table (Supporting Information 4) and individual diversity is represented using default parameters SPLITSTREE v4 (67; genes present in all individuals, 15.2 Mbp).

## Supporting information

Supporting Information 1 - Annotation of Uromyces beticola (beet rust)

Supporting Information 2 - Tables referenced in Supporting Information 1

Supporting Information 3 - Rust repeat annotation and comparison

Supporting Information 4 - Beet rust SNP diversity and divergence data

## Data availiblity

Illumina MP read data for the *U. beticola* genome and 46 re-sequenced individuals as well as 12 RNA-seq libraries used for annotation have been submitted to the European Nucleotide Archive (under projects xxx pending project numbers). The genome and annotation is available at the Earlham Institute Open Data site (pending xxx).

## Acknowledgments

This work was supported by the British Beet Research Organisation (BBRO) award (Discovering the source of sugar beet infection and re-infection) and the Biotechnology and Biological Sciences Research Council (BBSRC), part of UK Research and Innovation, through the Core Capability Grant (BB/CCG1720/1) and (BBS/E/T/000PR9818) WP1 Signatures of Domestication and Adaptation at the Earlham Institute (EI). We received funding for transfer of rust culturing skills via the BBSRC Flexible Talent Mobility Accounts (BB/R50659X/1). This work was delivered via the BBSRC National Capability in Genomics and Single Cell Analysis (BBS/E/T/000PR9816) at EI by members of the Genomics Pipelines and Core Bioinformatics Groups. Work was also delivered by Scientific Computing group and support for the physical HPC infrastructure and data centre delivered via the NBI Computing infrastructure for Science (CiS) group. HY was supported by the BBSRC funded Norwich Research Park Biosciences Doctoral Training Partnership grant BB/M011216/1 (project 2246665, with KWS UK Ltd (Student Project Partner). SW is supported by the Welcome Trust (218328/J/19/Z). We are also grateful to Wilfried Haerty and Conrad Nieduszynski (EI) for critical comments to the manuscript.

## Supporting information captions

**Supporting Information 1 – Annotation of Uromyces beticola (beet rust)**. Detailed methods and data organisation for the beet rust annotation.

**Supporting Information 2 – Tables referenced in Supporting Information 1**.

Genome – Assembly quality metrics such as N50

Reads Alignment – Mapping quality for the 12 RNA-Seq PE read libraries that were aligned to the genome

Transcript Assemblies – Transcript assembly metrics for two assembly methods

Repeats – Repeat masked regions for protein alignments and the gene build

Protein Alignments – Protein alignment summary from 26 representatives (including references)

Mikado Transcript – Mikado transcript assembly integration gene model statistics

Augustus Training – Augustus training results based on Mikado Gold transcripts

Augustus – Gene model statistics from three Augustus runs using different levels of evidence

Minos Release – Final gene model statistics after selection using Minos-Mikado

**Supporting Information 3 – Rust repeat annotation and comparison.** Reanalysis of repeat content of five published genomes (plus *U. beticola*).

**Supporting Information 4 – Beet rust SNP diversity and divergence data**.

