## Supporting Information 1 - Annotation of Uromyces beticola (beet rust) for "Analysis of wild plant pathogen populations reveals a signal of adaptation in genes evolving for survival in agriculture in the beet rust pathogen (*Uromyces beticola*)"

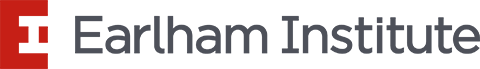

### Supporting Information 1 – Annotation of *Uromyces beticola* (beet rust)

Gemy Kaithakottil & David Swarbreck

Gene models were annotated using a workflow which incorporated repeat identification, RNA-Seq mapping and assembly, and alignment of protein sequences from related species. Alternative reference guided assembly methods were employed (Scallop, Kovaka et al., 2019; StringTie2, Shao and Kingsford, 2017)) to assemble transcripts for each sample. From these a filtered set of non-redundant transcripts were derived using Mikado (Venturini et al., 2018, see https://github.com/EI-CoreBioinformatics/mikado). Gene models were classified based on alignment to protein sequences, identifying the subset of gene models with likely full-length ORFs. The classified models together with aligned proteins and repeat annotation are provided as hints to AUGUSTUS (Stanke et al., 2006). Three alternative AUGUSTUS gene builds were generated using different evidence inputs or weightings. These together with the gene models derived from the Mikado transcript selection stage were consolidated into a single set of gene models using Minos (Venturini et al., 2018, https://github.com/EI-CoreBioinformatics/minos). The Minos pipeline scores alternative models based on the level of supporting evidence (protein homology, transcriptome data) and gene structure characteristics (e.g., CDS, UTR features) to select a representative gene model and alternative splice variants.

**EI core bioinformatics tools references:**

**Mikado -** [**https://github.com/EI-CoreBioinformatics/mikado**](https://github.com/EI-CoreBioinformatics/mikado)

Venturini, L., Caim, S., Kaithakottil, G. G., Mapleson, D. L., & Swarbreck, D. (2018). Leveraging multiple transcriptome assembly methods for improved gene structure annotation. GigaScience, 7(8). <https://doi.org/10.1093/gigascience/giy093>

**Portcullis -** [**https://portcullis.readthedocs.io/en/latest/**](https://portcullis.readthedocs.io/en/latest/)

Mapleson, D., Venturini, L., Kaithakottil, G., & Swarbreck, D. (2018). Efficient and accurate detection of splice junctions from RNA-seq with Portcullis. GigaScience, 7(12). <https://doi.org/10.1093/gigascience/giy131>

**Minos-Mikado – A gene model consolidation pipeline for genome annotation**

<https://github.com/EI-CoreBioinformatics/minos>

Final output files are provided within data package **UROBE1963355_EIv1.0_Frozen_release_10Jun2020.**

Final output files include gene positions and features (*gff3), FASTA files of extracted cDNA, CDS and protein sequences, summary files describing the annotated gene models and functional annotation (*tsv).

UROBE1963355_EIv1.0_Frozen_release_10Jun2020

└── EIv1.0

├── annotation

│	├── UROBE1963355_EIv1.0.annotation.gff3

│	├── UROBE1963355_EIv1.0.annotation.gff3.biotype_conf.summary

│	├── UROBE1963355_EIv1.0.annotation.gff3.cdna.fasta

│	├── UROBE1963355_EIv1.0.annotation.gff3.cds.fasta

│	├── UROBE1963355_EIv1.0.annotation.gff3.final_table.tsv

│	├── UROBE1963355_EIv1.0.annotation.gff3.metrics.tsv

│	├── UROBE1963355_EIv1.0.annotation.gff3.mikado_stats.txt

│	├── UROBE1963355_EIv1.0.annotation.gff3.mikado_stats.txt.summary

│	├── UROBE1963355_EIv1.0.annotation.gff3.pep.fasta

│	├── UROBE1963355_EIv1.0.annotation.gff3.pep.fasta.functional_annotation.tsv

│	└── UROBE1963355_EIv1.0.annotation.representative.gff3

└── assembly

├── Uromyces_beticola_EI_v1.1.genome.fasta

└── Uromyces_beticola_EI_v1.1.genome.fasta.fai

**Annotation statistics**

| Gene Model Classification |  |  |  |
| --- | --- | --- | --- |
| **Biotype** | **Confidence** | **Gene** | **Transcript** |
| **protein_coding_gene** | **High** | **6,954** | **14,908** |
| protein_coding_gene | Low | 928 | 1,224 |
| transposable_element_gene | High | 4,488 | 5,499 |
| transposable_element_gene | Low | 1,976 | 2,194 |
| predicted_gene | Low | 1,266 | 1,459 |
| Total | - | **15,612** | **25,284** |

| Gene Model Stats | All Genes | **All Genes (excluding transposable_element_gene)** |
| --- | --- | --- |
| Number of genes | 15,612 | **9,148** |
| Number of Transcripts | 25,284 | **17,591** |
| Transcripts per gene | 1.62 | **1.92** |
| Number of monoexonic genes | 1,590 | **322** |
| Monoexonic transcripts | 1,674 | **353** |
| Transcript mean size cDNA (bp) | 1,817.82 | **2,057.72** |
| Transcript median size cDNA (bp) | 1,570 | **1,790** |
| Min cDNA | 200 | **201** |
| Max cDNA | 16,251 | **16,251** |
| Total exons | 132,064 | **112,175** |
| Exons per transcript | 5.22 | **6.38** |
| Exon mean size (bp) | 348.03 | **322.69** |
| CDS mean size (bp) | 264.38 | **249.34** |
| Transcript mean size CDS (bp) | 1,116.92 | **1,330.55** |
| Transcript median size CDS (bp) | 814 | **1,020** |
| Min CDS | 71 | **71** |
| Max CDS | 15,627 | **15,627** |
| Intron mean size (bp) | 104.31 | **99.74** |
| 5UTR mean size (bp) | 283.99 | **285.85** |
| 3UTR mean size (bp) | 416.92 | **441.32** |

| **BUSCO v4.0 Stats (basidiomycota_odb10)** | **Genome** | **Transcripts** | **Proteins** |
| --- | --- | --- | --- |
| Complete (single copy) | 1,475 | 1,620 | 1,641 |
| Complete (2 copies) | 18 | 28 | 30 |
| Complete (3 copies) | 0 | 0 | 0 |
| Complete (4+ copies) | 0 | 0 | 0 |
| Complete | 1,493 | 1,648 | 1,671 |
| Duplicated | 18 | 28 | 30 |
| Fragmented | 24 | 18 | 18 |
| Missing | 247 | 98 | 75 |
| Total | 1,764 | 1,764 | 1,764 |

**Bioinformatics Processing**

**Summary**

1. Genome summary
2. RNA-seq alignment and transcriptome assembly
3. Gene predictor training
4. Cross-species protein alignment
5. Gene build
6. Gene model consolidation
7. Functional annotation

**Details:**

1. **Genome Summary**

Summary assembly statistics for **Uromyces_beticola_EI_v1.1.genome.fasta:**

| Mean: | 29,880.49 |
| --- | --- |
| Median: | 11,829 |
| Min: | 1,000 |
| Max: | 554,123 |
| N50[length]: | 294,203,835 |
| N50[value]: | 74,005 |
| L50: | 2,309 |
| Total_length: | 588,346,853 |
| Total_sequences: | 19,690 |

Please refer to the excel file (Supporting Information 2, tab **Genome**) for additional details.

2. **RNA-Seq alignment** **and transcript assembly**

The RNA-Seq PE reads were aligned to the genome using HISAT2 v2.1.0 (with option: --dta --min-intronlen=20 --max-intronlen=50000 --rna-strandness RF) - <https://daehwankimlab.github.io/hisat2/>. The aligned reads were assembled using StringTie2 v1.3.3 (with option: --rf) - <https://ccb.jhu.edu/software/stringtie/>, as well as Scallop v0.10.2 (with option: --library_type first) - <https://github.com/Kingsford-Group/scallop>. High confidence junctions were identified with Portcullis v1.1.2 (with options: full --threads 8 --orientation FR --canonical C,S --min_cov 2 --save_bad --strandedness firststrand) - <https://github.com/EI-CoreBioinformatics/portcullis>

Please refer to the excel file (Supporting Information 2, tabs **Reads_Alignment** and **Transcript_Assemblies**) for additional details.

Mikado was used to integrate the transcript assemblies (StringTie and Scallop) and select the best (highest scoring) model at each locus utilising intrinsic metrics based on ORF’s predicted using prodigal v2.6.3 (with options: -g 1 -f gff - <https://github.com/hyattpd/Prodigal>), and extrinsic metrics derived from BLASTX searches against the cross-species reference protein database using diamond v0.9.24 (with options: blastx --outfmt 6 qseqid sseqid pident length mismatch gapopen qstart qend sstart send evalue bitscore ppos btop - <http://www.diamondsearch.org/index.php>) and junctions passing portcullis filtering.

Please refer to the excel file (Supporting Information 2, tab **Mikado_Transcript**) for additional details.

3. **Gene predictor training**

Mikado primary ‘True’ transcript models were further classified into three categories, namely:

- Gold: Models with Full-lengtherNEXT (v0.0.8; Fernandez and Guerrero (2012); using fungal species downloaded on 04Feb2020) - <http://www.scbi.uma.es/site/scbi/downloads/313-full-lengthernext> hit of ‘Complete/Putative Complete’ and with at most two complete five_prime_UTR’s and three five_prime_UTR’s and at most one complete three_prime_UTR and two three_prime_UTR’s
- Silver: Models with CDS length >= 900bps and with at most two complete five_prime_UTR’s and three five_prime_UTR’s and at most one complete three_prime_UTR and two three_prime_UTR’s
- Bronze: Models that did not classify as Mikado transcript Gold or Silver

A subset of Mikado Gold transcripts were selected for training AUGUSTUS i.e. those with a single full length ORF, 5’ and 3’ UTR present, consistent Full-lengtherNEXT and CDS coordinates, a minimum CDS to transcript ratio of 50% and a single transcript per gene. We excluded genes with a genomic overlap within 1000bp of a second gene and gene models that are homologous to each other with a coverage and identify of 80%. The filtered Mikado Gold set contained 2412 transcripts for training AUGUSTUS. The trained AUGUSTUS model resulted in 0.963 sn, 0.915 sp nucleotide level, 0.732 sn, 0.718 sp exon level and 0.33 sn, 0.301 sp at the gene level.

Please refer to the excel file (Supporting Information 2, tab **Augustus_Training**) for additional details.

4. **Cross-species protein alignment**

The cross-species proteins were soft-masked using segmasker (blast v2.6.0) and were aligned to the low complexity soft-masked genome (RepeatMasker v4.0.7 Combined Database: Dfam_Consensus-20170127, RepBase-20170127 using pucciniaceae species) using exonerate v2.4.0 (with options: --model protein2genome --showtargetgff yes --showvulgar yes --querychunkid 10 -M 281.25 -D 281.25 --hspfilter 100 --softmaskquery yes --softmasktarget yes --bestn 10 --minintron 20 --maxintron 20000 --showalignment no --geneseed 250 --percent 30 --score 50 --ryo ">%qi\tlength=%ql\talnlen=%qal\tscore=%s\tpercentage=%pi\nTarget>%ti\tlength=%tl\talnlen=%tal\n") - <https://www.ebi.ac.uk/about/vertebrate-genomics/software/exonerate-manual> and the protein alignments were filtered at 50% identity and 80% coverage.

Please refer to the excel file (Supporting Information 2, tab **Protein_Alignments**) for more details.

5. **Gene build**

The evidence guided annotation of protein coding genes based on repeats, RNA-Seq mapping, transcript assembly and alignment of protein sequences was created using AUGUSTUS v3.3.3 (with options: --AUGUSTUS_CONFIG_PATH=trained_species_config –species= Uromyces_beticola --UTR=on --alternatives-from-evidence=true --noInFrameStop=true --allow_hinted_splicesites=atac) - <https://github.com/Gaius-Augustus/Augustus>.

**Repeats**

We used the RepeatModeler v1.0.10 - http://www.repeatmasker.org/RepeatModeler/, library of repeats from the earlier Uromyces_beticola_EI_v1.1.genome.fasta release of the genome.

Subsequent steps were:

1) RepeatMasker v4.0.7 - http://www.repeatmasker.org/ with RepBase Pucciniaceae library (RepBaseRepeatMaskerEdition-20170127.tar.gz)

2) RepeatMasker with the RepeatModeler library

And the above interspersed repeats were combined and used for the gene build.

Please refer to the excel file (Supporting Information 2, tab **Repeats**) for more details.

AUGUSTUS was run three different ways by assigning higher bonus scores and priority based on evidence type and classification (Gold, Silver, Bronze) to reflect the reliability of different evidence sets as described in the table below:

Run1: Run utilizes the evidence hints generated from Mikado transcript models, RNA-Seq Portcullis junctions, cross-species protein alignments (filtered at 80% coverage and 50% identity), RNA-Seq read coverage and interspersed repeats by using the sources and priorities described in the table below

Run2: Run uses same evidence hints as Run1, except that we do not use the RNA-Seq read coverage hints

Run3: Run uses same evidence hints as Run2, except that we give higher weightage to the cross-species protein alignments, and we also change the priorities as described in the table below

| Evidence | Run1 | Run2 | Run3 |
| --- | --- | --- | --- |
| Mikado Transcript Gold | Source M; Priority 10; | Source M; Priority 10; | Source E; Priority 10; |
| Mikado Transcript Silver | Source F; Priority 9; | Source F; Priority 9; | Source E; Priority 9; |
| Mikado Transcript Bronze | Source E; Priority 8; | Source E; Priority 8; | Source E; Priority 8; |
| Mikado Transcripts | Source E; Priority 7; | Source E; Priority 7; | Source E; Priority 7; |
| Portcullis Pass Gold  (score = 1) | Source E; Priority 6; | Source E; Priority 6; | Source E; Priority 6; |
| Portcullis Pass Silver  (score < 1) | Source E; Priority 4; | Source E; Priority 4; | Source E; Priority 4; |
| Proteins | Source P; Priority 4; | Source P; Priority 4; | Source P; Priority 9; |
| RNA-Seq Coverage Wig Hints | Source W; Priority 3; | NONE | NONE |
| Repeats | Source RM; Priority 1; | Source RM; Priority 1; | Source RM; Priority 1; |

Please refer to the excel file (Supporting Information 2, tab **Augustus**) for more details.

6. **Gene model consolidation**

The final set of gene models was selected using Minos-Mikado [**https://github.com/EI-CoreBioinformatics/minos**](https://github.com/EI-CoreBioinformatics/minos)**.** Minos is a pipeline that generates and utilises metrics derived from protein, transcript and expression data sets to consolidate gene models. In this annotation, the three alternative Augustus gene builds described earlier, and the gene models derived from the Mikado transcript run were consolidated into a single set of gene models.

Please refer to the excel file (Supporting Information 2, tab **Minos_Release**) for more details.

**Assignment of gene biotypes and confidence classification**

Gene models were classified as biotypes protein_coding_gene, predicted_gene and transposable_element_gene, and assigned as high or low confidence based on below criteria:

a) **High confidence protein_coding_gene:** Any protein coding gene where any of its associated gene models having a BUSCO v4.0.6 (Seppey et al., 2019) protein status of Complete/Duplicated OR having blastp (v2.9.0+) coverage (average across query and target coverage) >= 80% against the list protein datasets mentioned in Section 4. Cross-species protein alignments OR having average blastp coverage (across query and target coverage) >= 60% against the list protein datasets mentioned in Section 4. Cross-species protein alignments AND having transcript alignment F1 score (average across nucleotide, exon and junction F1 scores based on RNA-Seq transcript assemblies) >= 40%.
b) **Low confidence protein_coding_gene**: Any protein coding gene where all of its associated transcript models do not meet the criteria to be considered as high confidence protein coding transcripts.
c) **High confidence transposable_element_gene**: Any protein coding gene where any of its associated gene models having coverage >= 40% against the combined interspersed repeats mentioned in Section 5. Gene build under Repeats section.
d) **Low confidence transposable_element_gene**: Any protein coding gene where all of its associated transcript models do not meet the criteria to be considered as high confidence transposable_element_gene.
e) **Low confidence predicted_gene**: Any protein coding gene where any of its associated gene models having average blastp coverage (across query and target coverage) < 30% against the list protein datasets mentioned in Section 4. Cross-species protein alignments AND having a protein-coding potential score < 0.25 calculated using CPC2 0.1 (Kong et al., 2007)
f) Discarded models: Any models having no BUSCO protein hit AND no protein alignment score (average across nucleotide, exon and junction F1 scores based on protein alignments) AND no transcript alignment F1 score (average across nucleotide, exon and junction F1 scores based on RNA-Seq transcript assemblies) AND no blastp coverage (average across query and target coverage) AND Kallisto v0.44 (Bray et al., 2016) expression score <0.3 from across RNA-Seq reads OR having short CDS <30bps.
7. **Functional annotation**

All the proteins were annotated using AHRD v.3.3.3 (Hallab et al., 2014; https://github.com/groupschoof/AHRD/blob/master/README.textile). Sequences were blasted against the UniProt fungi sequences (data download date 04Jun2020), both Swiss-Prot and TrEMBL datasets (The UniProt Consortium, 2014). Proteins were BLASTed (v2.6.0; blastp) with an e-value of 1e-5. We have also provided InterProScan (v5.22.61; Jones et al., 2014) results to AHRD. We adapted the standard AHRD example configuration file path test/resources/ahrd_example_input_go_prediction.yml, distributed with the AHRD tool, changing the following apart from the location of input and output files:

1. we included the GOA mapping from uniprot (ftp://ftp.ebi.ac.uk/pub/databases/GO/goa/UNIPROT/goa_uniprot_all.gaf.gz) as parameter 'gene_ontology_result',

2. we also included the interpro database (ftp://ftp.ebi.ac.uk/pub/databases/interpro/61.0/interpro.xml.gz) and provided as parameter 'interpro_database',

3. we changed the parameter 'prefer_reference_with_go_annos' to 'false' and did not use the parameter 'gene_ontology_result',

4. we have only used swissprot and trembl as blast_dbs databases

A.F.A. Smit & R. Hubley RepeatModeler at <http://www.repeatmasker.org/RepeatModeler/>

A.F.A. Smit, R. Hubley & P. Green RepeatMasker at <http://repeatmasker.org>

Haas, B. J., 2010. TransposonPSI.

Kong, L., Zhang, Y., Ye, Z.-Q., Liu, X.-Q., Zhao, S.-Q., Wei, L., and Gao, G., 2007. CPC: assess the protein-coding potential of transcripts using sequence features and support vector machine. Nucleic Acids Research, 35(Web Server issue):W345–9. <https://doi.org/10.1093/nar/gkm391>

Bray, N. L., Pimentel, H., Melsted, P., and Pachter, L., 2016. Near-optimal probabilistic RNA-seq quantification. Nature Biotechnology, 34(5):525–527. <https://doi.org/10.1038/nbt.3519>

Hallab, A., Klee, K., Boecker, F., Girish, S., and Schoof, H., (2014). Automated assignment of Human Readable Descriptions (AHRD).

Apweiler, R. (2004). UniProt: the Universal Protein knowledgebase. Nucleic Acids Research, 32(90001), 115D – 119. <https://doi.org/10.1093/nar/gkh131>

Jones, P., Binns, D., Chang, H. Y., Fraser, M., Li, W., McAnulla, C., McWilliam, H., Maslen, J., Mitchell, A., Nuka, G., Pesseat, S., Quinn, A. F., Sangrador-Vegas, A., Scheremetjew, M., Yong, S. Y., Lopez, R., & Hunter, S. (2014). InterProScan 5: Genome-scale protein function classification. Bioinformatics, 30(9), 1236–1240. <https://doi.org/10.1093/bioinformatics/btu031>
