## Supporting Information 3 - Rust repeat annotation and comparison for "Analysis of wild plant pathogen populations reveals a signal of adaptation in genes evolving for survival in agriculture in the beet rust pathogen (*Uromyces beticola*)"

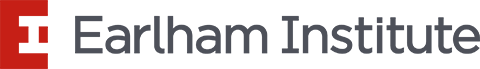

### Supporting Information 3 – Rust repeat annotation and comparison

Gemy Kaithakottil, Mark M^c^Mullan & David Swarbreck

#### Results & Discussion

The *Uromyces beticola* assembly is both large and has a high-level high repeat content. To identify whether the high levels of repeat content disproportionately affected this short read assembly we reanalysed and compared several rust assemblies. We find that the 588Mbp *U. beticola* assembly is of a similar size to *Hemileia vastatrix* (541Mbp) and yet *U. beticola* contains a lower proportion of missing and fragmented content (Fig. 1, see main text). Moreover, the *U. beticola* assembly contains levels of missing a duplicated content within the ranges of rust genomes that are between one third and one tenth the size (Fig. 1 main text; *Puccinia graminis* (81.5Mbp), *P. striiformis* f. sp. *Tritici* (61.4Mbp), *P. triticinia* (106.6Mbp) and *Uromyces viciae-fabae* (209.5Mbp)).

The *U. beticola* assembly contains a large proportion of interspersed repeat content (~90%). Reanalysis of repeat content in these rust genomes (using RepeatModeler RepeatMasker) shows a strong correlation between genome size and repeat content (Fig. 1). The evidence suggests that the expansion of the *U. beticola* and the proliferation of repeat content is in line with the level observed in other rusts.

Repeat content can facilitate the generation of structural genomic variation and this may be be particularly pertentent in this system where sexual reproduction and adaptation to wild and agricultural environments may impact on structural genomic variation. However, these processes cannot be rigorously investigated using this *U. beticola* short read assembly.

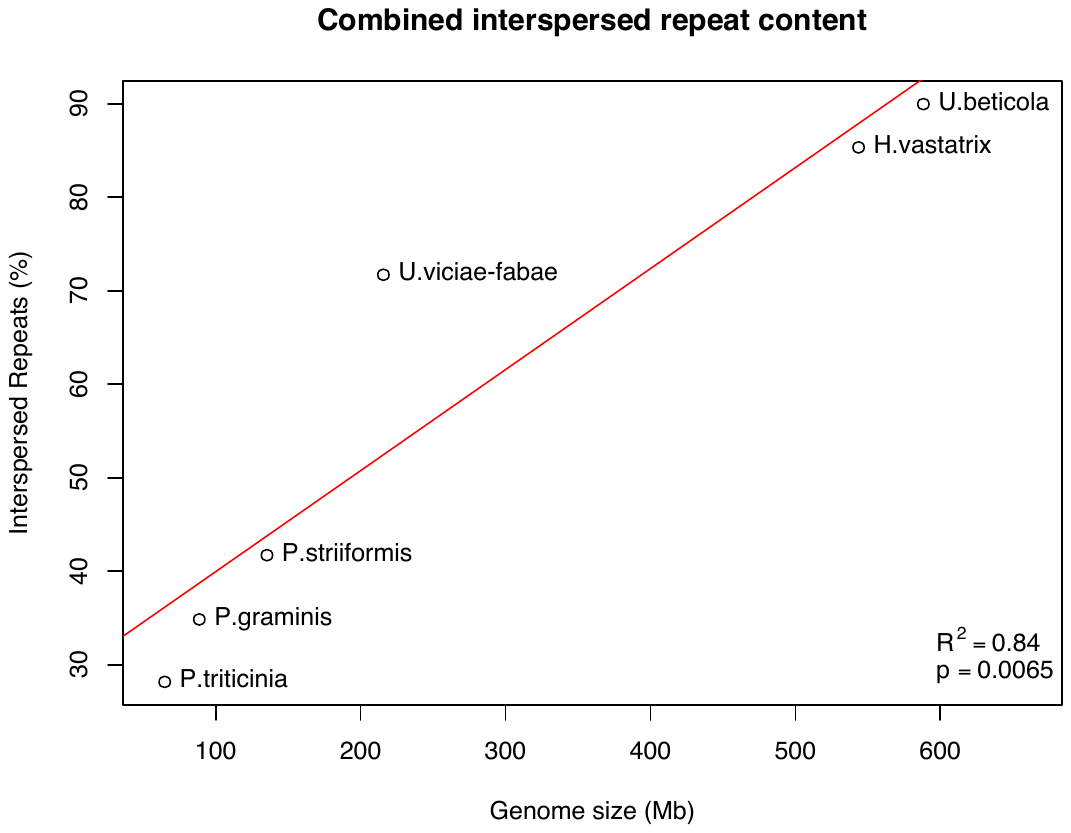

Figure 1. Combined interspersed repeat content relative to genome size. We observe a significant positive correlation between genome size and repeat content in the rust genomes analysed.

#### Methods

RepeatModeler v1.0.10 and RepeatMasker v4.0.7 were run for all assemblies using the methods described for the main *U. beticola* annotation (see Supporting Information 1).
